# Integrating vascular and hypertrophic cartilage microtissues to fabricate scaled-up grafts for endochondral bone tissue engineering

**DOI:** 10.64898/2026.07.13.738124

**Authors:** Gabriela S. Kronemberger, Ross Burdis, Catarina Correia, Leandra S. Baptista, Daniel J. Kelly

**Affiliations:** Trinity Centre for Biomedical Engineering, Trinity Biomedical Sciences Institute, Trinity College Dublin, Dublin, Ireland; Department of Mechanical, Manufacturing and Biomedical Engineering, School of Engineering, Trinity College Dublin, Dublin, Ireland; Department of Anatomy and Regenerative Medicine, Royal College of Surgeons in Ireland, Dublin, Ireland; Post-graduation Program of Translational Biomedicine (Biotrans), Unigranrio, Campus I, Duque de Caxias, Rio de Janeiro, Brazil; Nucleus of Multidisciplinary Research in Biology (Numpex-Bio), Federal University of Rio de Janeiro (UFRJ) Xerém, Duque de Caxias, Rio de Janeiro, Brazil; Advanced Materials and Bioengineering Research Centre (AMBER), Royal College of Surgeons in Ireland and Trinity College Dublin, Dublin, Ireland

## Abstract

The repair of large bone defects remains a major clinical challenge, in part due to inadequate vascularization and poor integration of graft materials. Tissue engineering strategies that recapitulate the developmental process of endochondral ossification, whereby a cartilage template remodels into bone, have shown significant potential in pre-clinical models of large bone defect healing. However, successfully scaling these approaches to clinically relevant sizes will require the development of strategies to support the rapid vascularization of the graft following implantation *in vivo*. Here, mechanically reinforced templates were first fabricated by integrating hypertrophic cartilage microtissues derived from human mesenchymal stem/stromal cells (MSCs) within an osteoconductive 3D-printed polycaprolactone (PCL) framework coated with nano-hydroxyapatite (nanoHA). *In vitro* the cartilage microtissues fused and generated an extracellular matrix rich in sulphated glycosaminoglycans and collagen. To prevascularize these constructs, vascular microtissues derived from a co-culture of endothelial cells and MSCs were incorporated into a central channel within the construct, which generated a microvascular network within the graft *in vitro*. Following subcutaneous implantation, hypertrophic cartilage templates with (‘vascular-channel’ group) and without (‘empty-channel’ group) this central vascularized channel supported endochondral bone formation. Quantitative microCT and histological analyses revealed significantly greater remaining bone in the empty-channel group, whereas the vascular-channel group supported enhanced vascularization and remodeling of the graft *in vivo*. These findings support the continued development and testing of a modular biofabrication strategy that combine self-organizing hypertrophic cartilage and vascular microtissues with osteoconductive 3D-printed architectures to generate scalable, prevascularised hypertrophic cartilage templates for endochondral bone repair.

**Key-words:** spheroids, microtissues, hypertrophic cartilage, vascularization, endochondral ossification, bone tissue engineering.

## 1. INTRODUCTION

Bone fractures arising from trauma, disease and ageing are amongst the most common injuries worldwide [1]. Although bone has an inherent capacity to heal, approximately 5–10% of fractures progress to delayed union or non-union and require surgical intervention [1–2]. The management of non-union remains a major clinical and socioeconomic burden, with reported treatment costs ranging from €10,000 to €100,000 per patient in Europe [3]. Autologous bone grafting remains the clinical gold standard. However, its use is constrained by donor-site morbidity, increased operative time and post-operative pain, and an elevated risk of complications, including procedure-related defect formation at the harvest site [4–5].

Endochondral ossification is a developmental process responsible for the formation of much of the mammalian skeleton, in which a cartilage template undergoes hypertrophy, vascular invasion and subsequent remodeling into bone through coordinated chondrocyte, vascular and osteoblast differentiation [6–7]. Unlike intramembranous ossification, which relies on the direct differentiation of mesenchymal stem cells (MSCs) into osteoblasts, the endochondral pathway is inherently adapted to survive in the low-oxygen, nutrient-poor environments characteristic of early-stage fracture healing [8]. Tissue engineering strategies that attempt to recapitulate key aspects of this process, often referred to as “developmental engineering”, represents a promising emerging treatment for large bone defects. By priming cells to form a hypertrophic cartilage intermediate *in vitro*, it is possible to engineer constructs that, upon implantation, initiate a cascade of host-mediated vascularization and remodeling into functional bone [9–10]. However, the clinical translation of such endochondral grafts remains constrained by the challenge of scaling these constructs to clinically relevant sizes. As construct dimensions increase, limited nutrient and oxygen diffusion restricts cell viability and tissue maturation *in vitro*, while insufficient or delayed vascularization following implantation compromises integration and functional bone formation *in vivo* [11–12].

The challenge of engineering tissues of clinically size can potentially be addressed by first fabricating numerous cellular aggregates, microtissues or organoids in isolation, can then combining these biological building blocks into larger regenerative templates [13–15]. Microtissues are typically generated by allowing cellular aggregates to self-assemble a tissue-specific extracellular matrix (ECM) *in vitro*. These modular units can then be combined using bottom-up bioassembly strategies to generate larger, clinically relevant constructs [16–17]. Importantly, this approach enables improved control over nutrient transport during *in vitro* culture while also providing an opportunity to incorporate specialized microtissue populations, such as vascular microtissues, to enhance perfusion and integration following implantation [18–19]. Several studies have explored MSC-derived microtissues for bone repair [20–24]. Fused microtissues can mineralize and express osteogenic markers, although fusion can be heterogeneous and structural organization is often limited without a supportive architecture [25]. More recently, microtissues have been combined with supporting polymeric scaffolds, which supported enhanced bone formation and functional repair in preclinical models [26]. Despite these advances, the challenges of scaling-up remains and most endochondral microtissue-based strategies have not considered establishing a functional vascular network within the construct prior to implantation.

Vascular microtissue-based strategies have also been explored to enhance vascularization and the osteogenic potential of engineered grafts [19;27]. For example, multicellular vascular spheroids combined with osteogenic cell sources within matrix-based bioinks have been associated with increased osteogenic and angiogenic gene expression and improved bone formation and neovascularization *in vivo* [28]. This raises the possibility that vascular microtissues could be combined with hypertrophic cartilage microtissues to prevascularize endochondral constructs, thereby overcoming a key barrier to their clinical translation. We previously reported a prevascularization strategy for hypertrophic cartilage microtissues using a fibrin-based hydrogel containing HUVECs, which enhanced mineralization and increased bone formation after subcutaneous implantation [29]. However, the advanced maturation state of the microtissues limited their fusion within the constructs, potentially limiting homogenous bone development and functional integration *in vivo*.

Therefore, the aim of this study was to biofabricate prevascularised human hypertrophic cartilage templates using microtissues as modular building blocks and to evaluate their capacity to support vascularization and endochondral bone formation *in vivo*. Hypertrophic cartilage microtissues were integrated within an osteoconductive, nanoHA-coated 3D-printed polycaprolactone (PCL) scaffold [30] incorporating a central channel that was fabricated using vascular microtissues engineered using a MSC–HUVEC co-culture. Following *in vitro* characterization, these constructs were implanted subcutaneously into nude mice to determine whether this prevascularization strategy enhances neovascularization and bone formation relative to non-pre-vascularized controls.

## 2. MATERIAL AND METHODS

### 2.1 Isolation and expansion of human bone marrow mesenchymal stem cells (bmMSCs)

Human bmMSCs were isolated from unprocessed human bone marrow aspirate (Lonza) on the basis of plastic adherence. Briefly, unprocessed bone marrow was plated at a density of 2.5×10^5^ cells/cm^2^ in expansion medium (X-Pan) composed of: Dulbecco’s Modified Eagle Medium (DMEM) + 100 U/mL penicillin + 100 μg/mL streptomycin (Gibco) + 10% (w/v) Fetal Bovine Serum (FBS) + 5 ng/mL Basic Fibroblast Growth Factor 2 (FGF-2) and cultured under physioxic conditions (37◦C in a humidified atmosphere with 5% CO_2_ and 5% O_2_). Following colony formation at 80% confluency, human bmMSCs were trypsinized using 0.25% (w/v) Trypsin Ethylenediaminetetraacetic acid (EDTA) and aggregated into microtissues at passage 3 (P3).

### 2.2 Human Umbilical Vein Endothelial cells (HUVECs) culture and expansion

GFP-transfected HUVECs (Lonza, Walkersville, MD) were cultured at 2,500 cells/cm^2^ in Endothelial Growth Medium (EGM-2) which had been supplemented with EGM-2 BulletKit (Lonza) following company recommendations and cultured at 37°C in a humidified atmosphere with 5 % CO_2_ and 20 % O_2_. HUVECs were aggregated into microtissues at passage 4 (P4).

### 2.3 Hypertrophic cartilage microtissue production

Human bmMSC derived microtissues were fabricated using an in-house medium high-throughput non-adherent agarose hydrogel microwell system as described previously (Nulty et al., 2021). In this system, it was possible to fabricate up to 401 microtissues at once. For microtissue formation, MSCs at passage 3 were seeded into each mould at a final density of 4 x 10^3^ cells/microtissue in X-Pan media. After 24h to allow for cell aggregation, the microtissues were induced using a chondrogenic defined media (CDM) consisting of hgDMEM GlutaMAX supplemented with 100 U/mL penicillin, 100 μg/mL streptomycin (both Gibco), 100 μg/mL sodium pyruvate, 40 μg/mL L-proline, 50 μg/mL L-ascorbic acid-2-phosphate, 4.7 μg mL/1 linoleic acid, 1.5 mg/mL bovine serum albumin, 1 × insulin–transferrin–selenium, 100 nM dexamethasone (all from Sigma), and 10 ng/mL of human transforming growth factor-β3 (TGF-β3) (Peprotech, United Kingdom) at 37°C in a humidified atmosphere with 5 % CO_2_ and 5 % pO_2_ for 2 days. The constructs were then moved to 20 % pO_2_ and switched to hypertrophic media consisting of hgDMEM GlutaMAX supplemented with 100 U/mL penicillin, 100 μg/mL streptomycin (both Gibco), 1 × insulin–transferrin–selenium, 4.7 μg/mL linoleic acid, 50 nM thyroxine, 100 nM dexamethasone, 250 μM ascorbic acid and 7 mM β-glycerophosphate (all from Sigma) for 1 week.

### 2.4 HUVEC microtissues production

For the generation of vascular spheroids, HUVECs (either alone or in combination with hMSCs in the ratios of 3MSC:1HUVEC; 1MSC:1HUVEC and 1MSC:3HUVEC) were seeded into the microwells as a homogenous cell suspension at an appropriate concentration to create spheroids contain 1 × 10^3^ cells. Microwells were centrifuged at 600 × g for 5 minutes and plates were then incubated in EGM, described previously, in physiological oxygen conditions (37 °C in a humidified atmosphere with 5 % CO2 and 5 % pO2) for 24 h to allow for spheroid formation.

### 2.5 Fabrication of fibrin-gelatin-hyaluronic acid hydrogels

Briefly, hyaluronic acid (Sigma) was added to high glucose Dulbecco’s Modified Eagle Medium (hgDMEM) (Gibco, Biosciences) at a concentration of 3 mg/mL and stirred overnight at 37°C. Next, 10% (v/v) glycerol (Sigma) was added, and the solution was stirred for 1 h at room temperature. Gelatin type A (175 g bloom) (Sigma) was added at a concentration of 40 mg/mL and stirred for 2 h at 37°C until fully dissolved. Before use, Fibrinogen (Sigma) was added to this thawed carrier gel at a concentration of 30 mg/mL and stirred for 2 h at 37°C.

The capacity to form functional vascular spheroids was evaluated using a fibrin-based hydrogel. Vascular spheroids were seeded into the hydrogels at a density of 300 spheroids per 50 μL of hydrogel. To enhance vascularization, EGM was supplemented with 50 ng/mL of human vascular endothelial growth factor (hVEGF, Peprotech). After 7 days of culture in physiological oxygen conditions (37 °C in a humidified atmosphere with 5 % CO2 and 5 % pO2), prevascularised constructs were evaluated using confocal microscopy.

### 2.6 Biofabrication of prevascularised constructs

A final number of 700 cartilage primed microtissues at 2 days maturation level were manually seeded in cylindrical 3D printed polycaprolactone (PCL) scaffolds. Briefly, scaffolds were designed using BioCADTM *software* package and produced using the 3D Discovery multi-head bioprinting system (RegenHu, Switzerland). Porous PCL (CAPA Ingevity, SC, United States) disc scaffolds were printed at 80°C at 0.5 MPa using a 27G needle (1 mm height, 2 mm diameter). Scaffolds were sterilized using ethylene oxide (EtO) gas prior to the addition of the microtissues. To assist the accuracy of the printing process, 4 % (w/v) agarose (Sigma) well inserts were fabricated using custom-designed moulds printed with Form2 SLA printer (FormLabs, MA, United States). A total of 300 3:1 MSC: HUVEC derived microtissues encapsulated in fibrin-gelatin-hyaluronic acid hydrogels were seeded in the central channel of the PCL groups. All constructs were then cultured in EGM-2 media supplemented with 50 ng/mL of VEGF for 7 days under normoxic conditions to allow for the formation of microvessels.

### 2.7 Biochemical analysis

Samples were washed in PBS after retrieval, and the wet weight was recorded for samples which were cultured dynamically. A papain enzyme solution, 3.88 U/mL of papain enzyme in 100 mM sodium phosphate buffer/5 mM Na2EDTA/10 mM Lcysteine, pH 6.5 (all from Sigma–Aldrich), was used to digest the samples at 60 °C for 18 hours. DNA content was quantified immediately after digestion using Quant-iT™ PicoGreen ® dsDNA Reagent and Kit (Molecular Probes, Biosciences). The amount of sGAG was determined using the dimethylmethylene blue dye-binding assay (Blyscan, Biocolor Ltd., Northern Ireland), with a chondroitin sulphate standard read using the Synergy HT multi-detection micro-plate reader (BioTek Instruments, Inc) with a wavelength set to 656 nm. Total collagen content was determined using a chloramine-T assay to measure the hydroxyproline content and calculated collagen content using a hydroxyproline-to-collagen ratio of 1:7.69. Briefly, samples were mixed with 38 % HCL (Sigma) and incubated at 110 °C for 18 hours to allow hydrolysis to occur. Samples were subsequently dried in a fume hood and the sediment reconstituted in ultra-pure H2O. 2.82 % (w/v) Chloramine T and 0.05 % (w/v) 4-(Dimethylamino) benzaldehyde (both Sigma) were added and the hydroxyproline content quantified with a trans-4-Hydroxy-L-proline (Fluka analytical) standard using a Synergy HT multi-detection micro-plate reader at a wavelength of 570 nm (BioTek Instruments, Inc).

### 2.8 Immunocytochemical analysis

Hydrogels were fixed with 4 % PFA for 30 minutes, permeabilized in 0.5 % Triton in PBS for 15 minutes and stained with Rhodamine red Phalloidin (5U/mL; 165nM; Thermo Fisher) for 1 h. Cells were washed and counterstained with DAPI (4′,6-diamidino-2-phenylindole; 1μg/mL; Sigma) for 15 minutes and imaged using a Leica SP8 scanning confocal microscope. Representative images are presented as maximum projections of z-stacks.

### 2.9 In vivo subcutaneous implantation

To investigate the effect of pre-vascularization on a hypertrophic cartilage template *in vivo*, three groups were compared: empty PCL (control), hypertrophic-cartilage and vascularized hypertrophic-cartilage were implanted subcutaneously into nude mice and explanted at 8 weeks for analyses. Constructs were implanted subcutaneously into the back of Balb/c female nude mice (Harlan, United Kingdom). Briefly, two subcutaneous pockets were made along the central line of the spine, one at the shoulders and the other at the hips. Two constructs were inserted into each pocket. Six constructs were implanted per group, and constructs were harvested at 8 weeks post-implantation. Mice were anaesthetized using an intraperitoneal injection of xylazine hydrochloride and ketamine hydrochloride, Carprofen was added to water for 24 h post-surgery, and mice were sacrificed by CO_2_ inhalation. This protocol and study were reviewed and approved by the ethics committee of Trinity College Dublin and the Health Products Regulatory Agency (HPRA, approval number AE19136/P069). Post-implantation samples were fixed in 4% PFA for 24 h.

### 2.10 Micro-computed tomography

Micro-computed tomography (μCT) scans were performed using a Scanco Medical 40 μCT system (Scanco Medical, Bassersdorf, Switzerland) to visualize and quantify mineral content and to assess mineral distribution within all constructs. Constructs were scanned in 50% EtOH, at a voxel resolution of 12 μm, a voltage of 70 kVp, a current of 114 μA. Reconstructed 3D images were generated to visualize the mineral content and distribution throughout the constructs. A Gaussian filter (sigma = 1.2, support = 2) was used to suppress noise and a global threshold of 482.

### 2.11 Histological analysis

*In vitro* samples were fixed using 4 % paraformaldehyde (PFA) solution overnight at 4 °C. After fixation, samples were dehydrated in a graded series of ethanol solutions (70 % - 100 %), cleared in xylene, and embedded in paraffin wax (all Sigma-Alrich). Prior to staining tissue sections (5 μm) were rehydrated. Sections were stained with hematoxylin and eosin (H&E), 1 % (w/v) alcian blue 8GX in 0.1 M hydrochloric acid (HCL) (AB) to visualize sulphated glycosaminoglycan (sGAG) content and counter-stained with 0.1 % (w/v) nuclear fast red to determine cellular distribution, and 0.1 % (w/v) picrosirius red (PSR) to visualize collagen deposition (all from Sigma-Aldrich). Stained sections were imaged using an Aperio ScanScope slide scanner.

*In vivo* samples were decalcified using “Decalcifying Solution-Lite” (Sigma) for approximately 1 month. *In vivo* samples were frequently x-rayed to determine if any mineral content remained. When no mineral was visible the sample was considered decalcified. All samples (*in vitro* and decalcified *in vivo*) were then dehydrated in graded series of ethanol solutions (70–100%), cleared in xylene, and embedded in paraffin wax (all Sigma-Aldrich). Sections (5-7 μm) were rehydrated in graded series and stained with Hematoxylin and eosin (H&E), 1 % (w/v) alcian blue (AB) 8GX in 0.1 M HCL to assess sulphated glycosaminoglycan (sGAG) content with a counter stain of 0.1% (w/v) nuclear fast red to assess cellular distribution, 0.1% (w/v) picrosirius red (PSR) to assess collagen distribution, 0.2% (w/v) Safranin O (Saf-O) to assess sGAG content post-implantation and Goldner’s trichrome (GT, Groat’s iron Hematoxylin, Fuchsine, Orange G, Fast Green; all from Sigma) for visualizing bone (Mineralized bone = dark green, Fibrous Tissue = Light green, Osteoid = orange/red, erythrocytes = dark red). Slides were then imaged using an Aperio ScanScope slide scanner. It should be noted that PCL is cleared during the tissue processing and leaves empty spaces in constructs as a result. *In vivo* sections were evaluated for vessel infiltration by counting vessels visible across an entire section using Aperio ImageScope and ImageJ software.

### 2.12 Image quantification analysis

#### 2.12.1 Masson’s Trichrome

Stained histological sections were quantitatively analyzed using a custom Python pipeline (OpenCV, NumPy). Whole-slide .tif images were imported and down sampled to 75% of their original linear resolution using area-based interpolation (cv2.INTER_AREA) to reduce computational cost while preserving colour fidelity. Images were converted from BGR to HSV colour space (cv2.cvtColor) to enable separation of chromatic information (hue and saturation) from intensity (value). Tissue components were segmented into three colour classes using empirically defined HSV thresholds: dark green (bone-associated staining; H: 35–130, S: 150–255, V: 50–175), light green (non-bone/connective tissue; H: 35–130, S: 20–180, V: 176–255), and dark red (muscle/erythrocytes; H: 130–255, S: 30–255, V: 40–200). Binary masks were generated using cv2.inRange, and segmentation performance was qualitatively verified by overlaying masks onto the original images (cv2.bitwise_and). For each class, positive pixels were quantified (cv2.countNonZero) and expressed as a fraction of the total image area to estimate relative tissue abundance. To account for differences in overall staining extent, these values were normalized to the total segmented signal (sum of all classes), yielding the proportional contribution of each tissue type within stained regions. Staining intensity was assessed independently of area by analyzing the saturation (S) channel. Masked saturation values were extracted for each class, and the mean saturation was calculated (cv2.mean) as a measure of colour intensity within each tissue compartment.

#### 2.12.2 Safranin O

Images were analyzed using a custom Python pipeline (OpenCV). Images were imported in BGR format and down sampled to 50% of their original linear resolution using area-based interpolation (cv2.INTER_AREA) to reduce computational cost while preserving colour information. Images were converted to HSV color space (cv2.cvtColor) to separate chromatic features (hue and saturation) from intensity (value), improving the robustness of colour-based segmentation. Two color classes were defined using empirically derived HSV thresholds: blue-stained regions (bone-associated/counterstain; H: 90–128, S: 50–255, V: 70–255) and red-stained regions (cartilage-associated Safranin O signal; H: 129–180, S: 50–255, V: 70–255). Binary masks were generated using cv2.inRange, and segmentation performance was qualitatively verified by overlaying masks onto the original images (cv2.bitwise_and). For each class, positive pixels were quantified (cv2.countNonZero) and expressed as a fraction of total image area to estimate relative staining abundance. To account for background and non-stained regions, class fractions were further normalized to the combined segmented signal (blue + red), yielding the relative composition of the stain-positive regions. Staining intensity was evaluated independently of area by analyzing the saturation (S) channel. Masked saturation values were extracted for each class, and the mean saturation was calculated (cv2.mean) as a measure of stain intensity for blue- and red-classified regions.

### 2.13 Statistical analysis

Statistical analyses were performed using GraphPad Prism software (GraphPad Software, CA, United States). To analyze significant differences between two groups at one timepoint a standard two-tailed t-test was performed. To analyze variance between > 2 groups at one timepoint one-way analysis of variance (ANOVA) was performed with Tukey post hoc test. To analyze variance between > 2 groups at multiple timepoints, *two-way* ANOVA was used with Tukey post hoc test. Numerical and graphical results are displayed as mean ± standard deviation unless stated otherwise. Significance was accepted at a level of p < 0.05.

## 3. RESULTS

### 3.1 Biofabrication of hypertrophic cartilage within 3D printed PCL scaffolds

Following 2 days of chondrogenic priming, human MSC derived microtissues were seeded into an nHA-coated PCL framework and cultured under dynamic conditions for 3 weeks (2 weeks of chondrogenic priming followed by 1 week of hypertrophic priming; Fig. 1). Histological analysis demonstrated cohesive fusion of the microtissues, with no discernible microtissue remnants observed within the construct (Fig. 1A). The extracellular matrix stained strongly for sulphated glycosaminoglycan (sGAG) deposition, as evidenced by intense Alcian Blue staining. Collagen and calcium deposition were preferentially localized to the peripheral regions of the constructs, as revealed by Picrosirius Red and Alizarin Red staining, respectively (Fig. 1A). These histological findings were supported by biochemical analyses, which revealed sGAG and collagen contents of approximately 60 µg/µg DNA and 50 µg/µg DNA, respectively (Fig. 1B).

**Figure 1.**
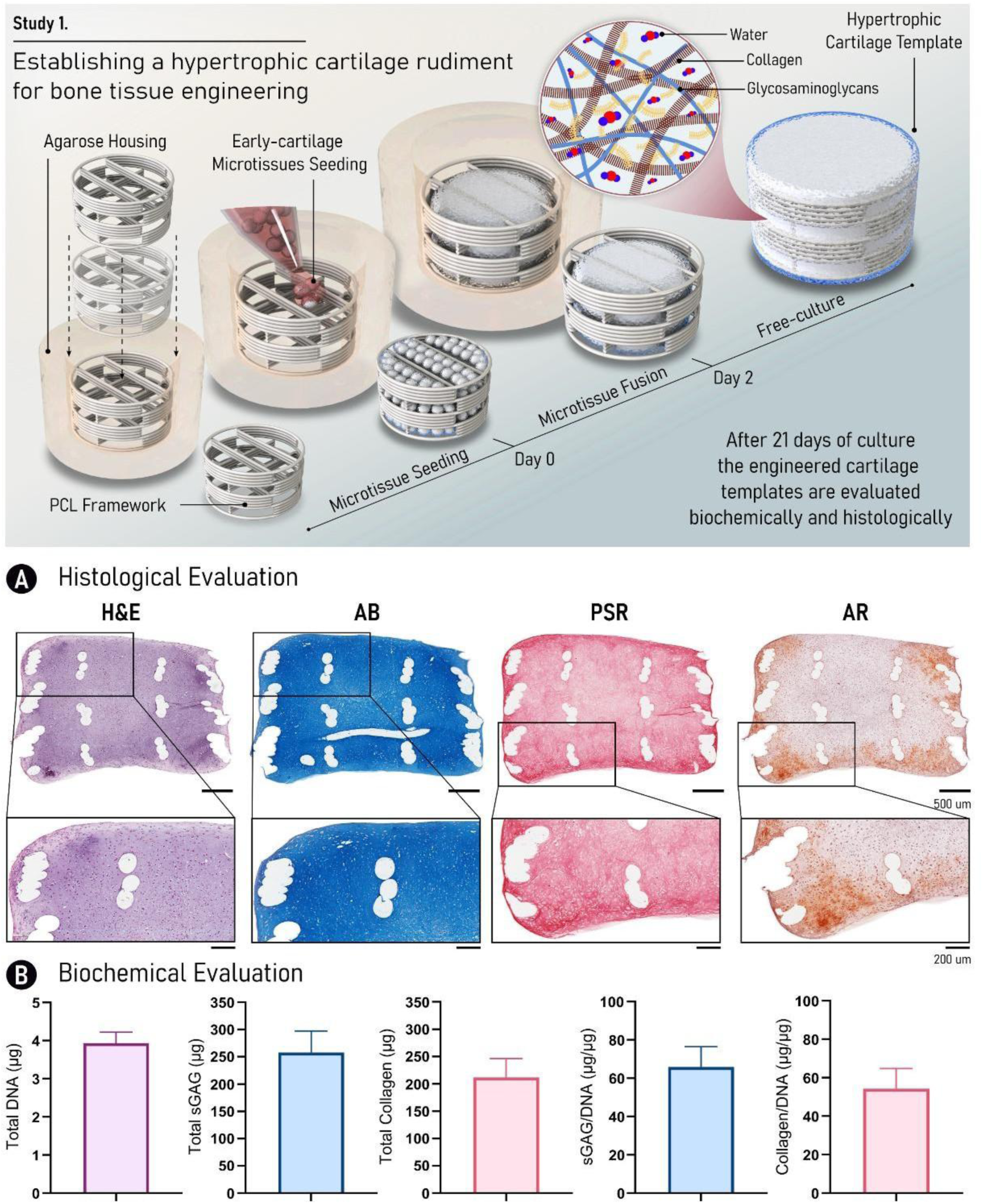
Hypertrophic cartilage templates exhibit cohesive microtissue fusion and robust extracellular matrix deposition following dynamic culture. Schematic overview of the study design is shown. (A) Histological evaluation at week 3 of culture using Hematoxylin and Eosin (H&E), Alcian Blue (AB), Picrosirius Red (PSR), and Alizarin Red (AR) staining. (B) Biochemical quantification of DNA, sulphated glycosaminoglycans (sGAG), collagen, and corresponding sGAG/DNA and collagen/DNA ratios at week 3 of culture. Scale bar: 200 µm.

### 3.2 The integration of vascular microtissues into engineered hypertrophic cartilage constructs

The engineering of vasculature remains a central challenge in the field of bone tissue engineering, which could potentially be addressed by the integration of vascular microtissues into a microtissue based grafts. Therefore, we next sought to biofabricate vascular microtissues by using different ratios of human bone marrow derived MSCs and HUVECs (Fig. 2A). All microtissues were homogeneous in size and shape, as observed by phase contrast images and confirmed by hematoxylin and eosin histological staining (Fig. 2B). After 24 hours, co-cultured and HUVEC-only microtissues were harvested and encapsulated in fibrin hydrogels for 7 days to evaluate micro-vessel formation *in vitro* (Fig. 2C). Fluorescence analyses revealed that the 3:1 MSC:HUVEC microtissue group supported more evident sprouting and microvessel formation (Fig. 2C), followed by the 1:1 MSC:HUVEC group (Fig. 2C). No sprouting or microvessel formation was observed in the 1:3 MSC:HUVEC and HUVEC only groups (Fig. 2C). This data suggests that vascular microtissues, when supported with appropriate numbers of MSCs, have the potential to support microvessel network development *in vitro*.

**Figure 2.**
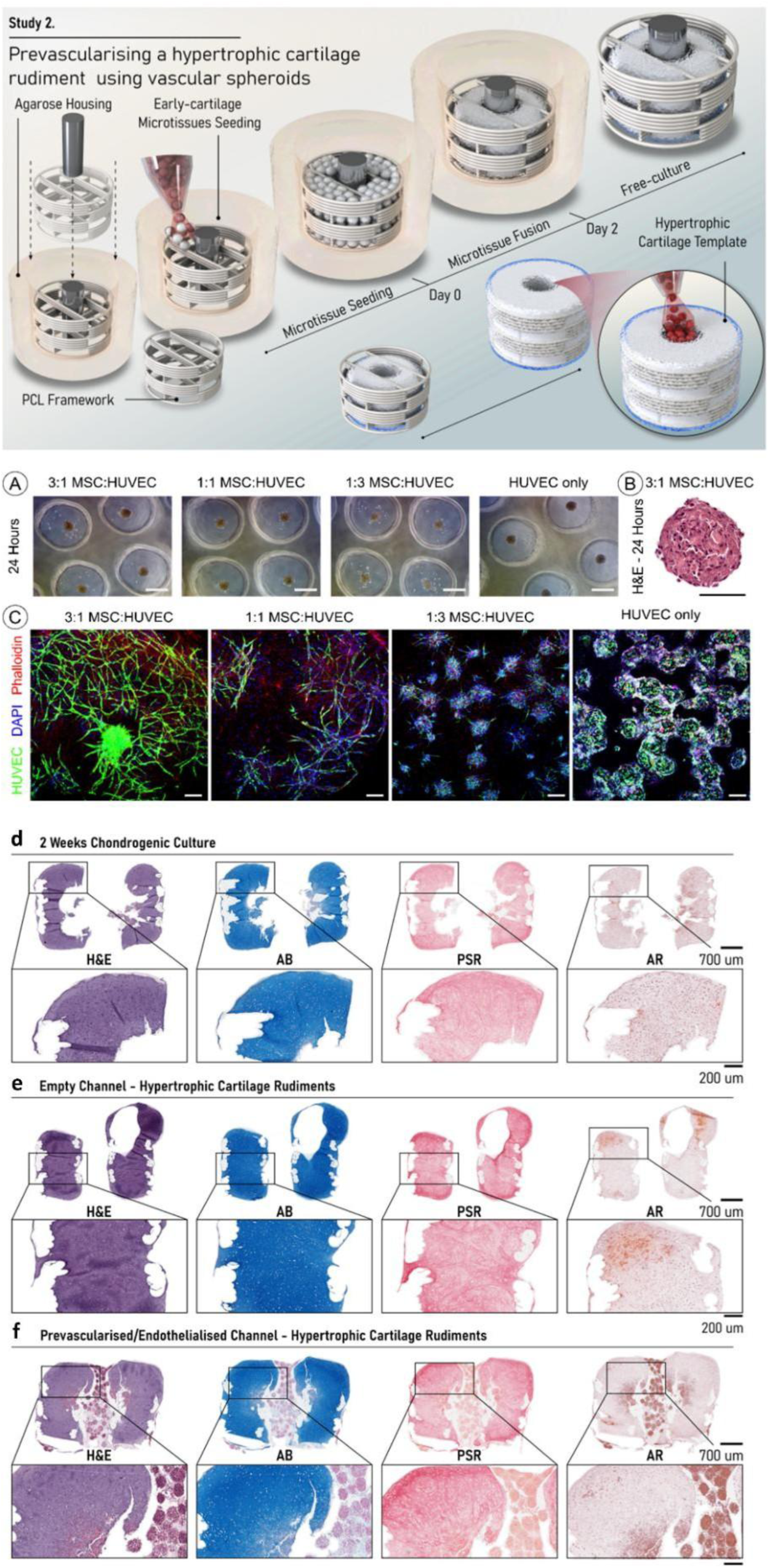
Biofabrication and characterization of vascular microtissues and vascularized hypertrophic cartilage templates. Schematic overview of the study design is shown. (A) Phase-contrast images of MSC:HUVEC co-cultures at different ratios, as well as HUVEC-only microtissues, used for vascular microtissue biofabrication. (B) Hematoxylin and eosin (H&E) staining of 3:1 MSC:HUVEC vascular microtissues after 24 h of culture. (C) Fluorescence imaging of co-culture and HUVEC-only vascular microtissues encapsulated in fibrin hydrogel after 1 week of culture. Samples were stained with phalloidin (red) for actin, DAPI (blue) for nuclei, and GFP (green) for HUVECs. Scale bar: 100 µm. Histological evaluation using H&E, Alcian Blue (AB), Picrosirius Red (PSR), and Alizarin Red (AR) staining of (D) cartilage templates after 2 weeks of culture, (E) hypertrophic cartilage templates after 3 weeks of culture, and (F) vascularized hypertrophic cartilage templates after 4 weeks of culture. Scale bars: 200 µm and 700 µm.

With the ultimate goal to biofabricate a vascularized-hypertrophic cartilage template, the 3:1 MSC:HUVEC vascular microtissues, encapsulated within a fibrin hydrogel, were assembled into a channel within the center of the hypertrophic-cartilage construct and cultured for the final week of a 4-week culture (2 weeks of chondrogenic priming, 1 week of hypertrophic priming and 1 week of vascular priming, Fig. 2 D-F). Histological assessment confirmed calcium deposition within the hypertrophic cartilage template, which was not observed following the chondrogenic priming phase (Fig. 2D-E). Histological assessment also confirmed the presence of the vascular microtissues in the central channel of the hypertrophic-cartilage template at the end of the culture period (Fig. 2F). However, the vascular microtissues did not fuse after seeding in the PCL scaffold, which might be explained by the presence of the fibrin gel (Fig. 2F). The extracellular matrix outside of the vascularized channel was rich in sGAG (AB) and collagen (PSR).

### 3.3 Pre-vascularization of hypertrophic cartilage templates increases vessel formation and accelerates cartilage remodeling in vivo

The potential of the different hypertrophic cartilage templates to produce new bone was next investigated in a subcutaneous nude mouse model (Fig. 3). Macroscopically it was observed that both experimental groups (empty channel’ and ‘vascular channel’) supported a higher amount of new tissue formation when compared to the empty PCL control group (Fig. 3A). In addition, the vascular microtissues were still observed in the central channel of the vascular channel group after the implantation period (Fig. 3A). MicroCT analysis revealed no bone formation in the central channel for both experimental groups (Fig. 3B), with significantly higher (p < 0.05) new bone formation observed in the empty channel hypertrophic cartilage construct compared to the pre-vascularized template (Fig. 3C). In addition, no bone formation was identified in the empty PCL group (control).

**Figure 3.**
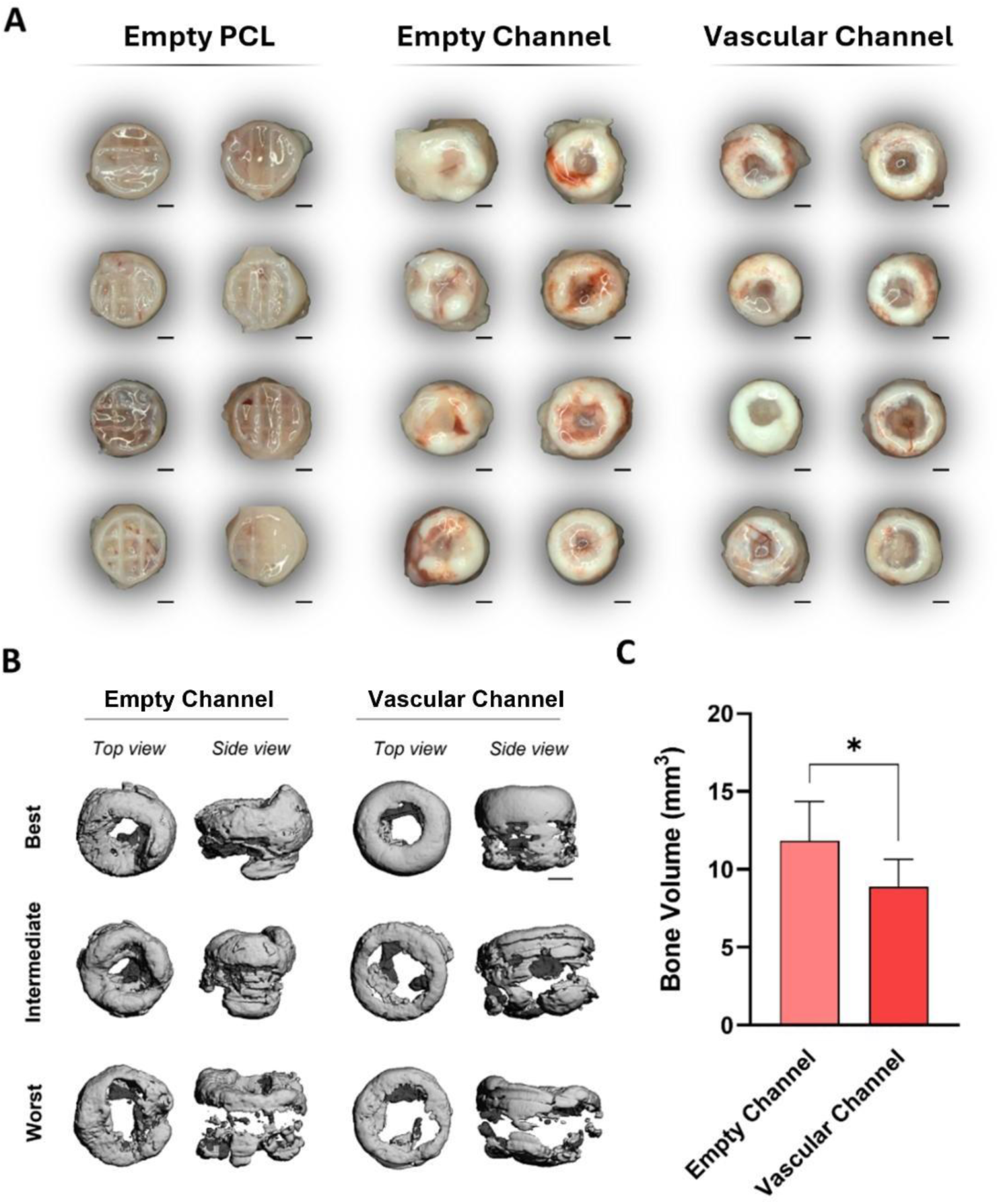
Hypertrophic cartilage templates exhibit higher bone volume following subcutaneous implantation. (A) Macroscopic images of empty PCL scaffolds (control), hypertrophic cartilage templates (empty channel), and vascularized hypertrophic cartilage templates (vascular channel) after 8 weeks of subcutaneous implantation in nude mice. (B) Representative microCT images showing best, intermediate, and worst samples from hypertrophic cartilage and vascularized hypertrophic cartilage template groups. (C) Quantification of bone volume based on microCT analysis of hypertrophic cartilage and vascularized hypertrophic cartilage templates. Data are presented as mean ± SD. Statistical significance was determined using an unpaired Student’s t-test (p < 0.05). Scale bar: 1 mm.

Histological analyses revealed pronounced differences between the empty channel and vascular channel hypertrophic cartilage templates following 8 weeks *in vivo* (Fig. 4A–C). Safranin O staining demonstrated higher levels of sGAG positive, residual cartilage tissue in the empty channel compared to the vascular channel templates (Fig. 4B), with quantification revealing significantly increased pixel intensity in the empty channel group (p < 0.0001; Fig. 4D). This suggests that pre-vascularization was accelerating the remodeling of the hypertrophic cartilage graft in vivo.

**Figure 4.**
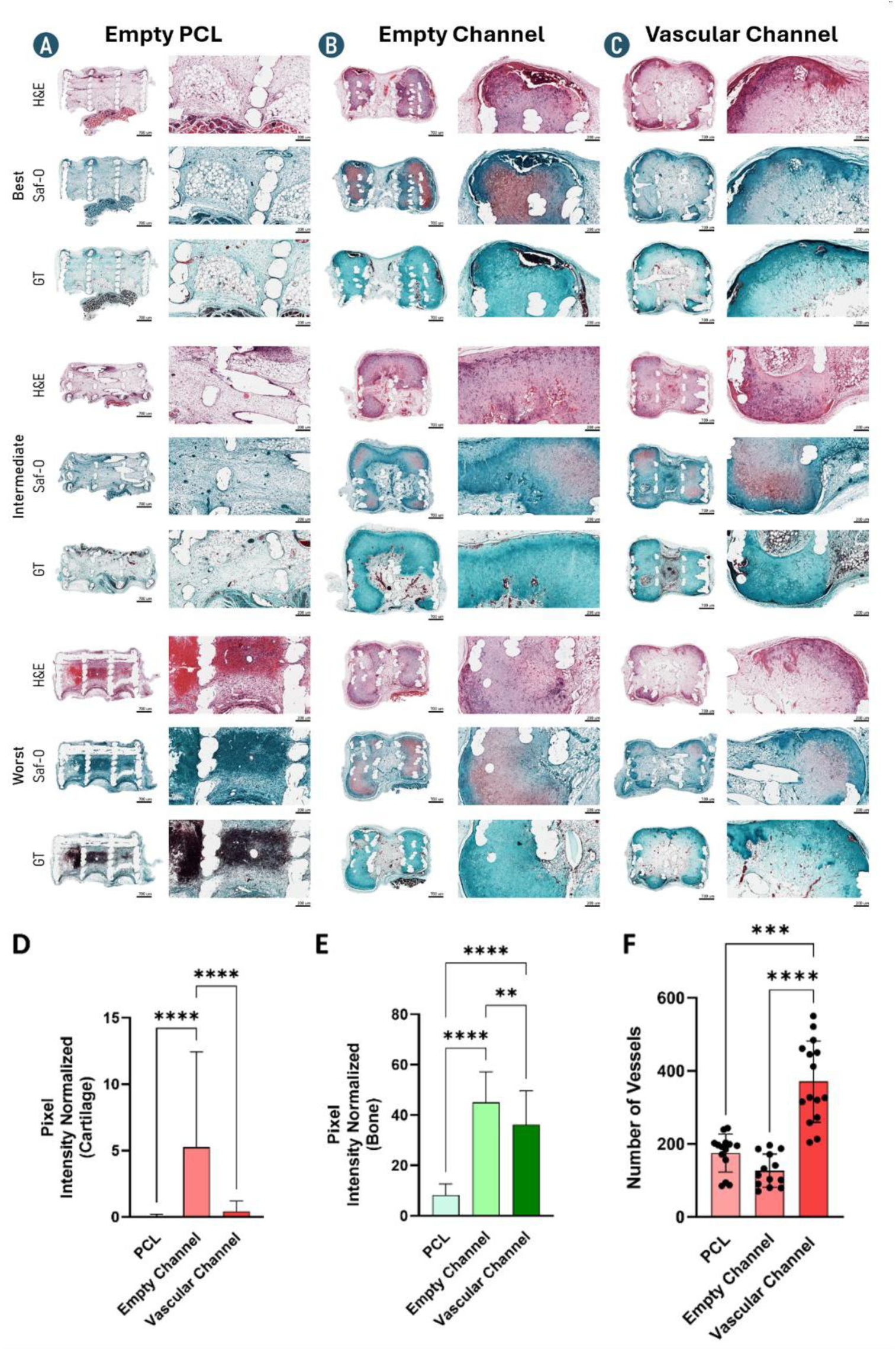
Hypertrophic cartilage templates exhibit higher sGAG deposition, whereas vascularized templates show increased microvessel formation. Histological evaluation using Hematoxylin and Eosin (H&E), Safranin O (Saf-O), and Goldner trichrome (GT) staining of representative best, intermediate, and worst samples from (A) empty PCL scaffolds (control), (B) hypertrophic cartilage templates (empty channel), and (C) vascularized hypertrophic cartilage templates. (D) Quantification of cartilage formation based on Safranin O pixel intensity. (E) Quantification of mature bone formation based on Goldner trichrome pixel intensity. (F) Quantification of vessel number. Data is presented as mean ± SD. The best, intermediate and worst samples were defined based on the levels of new bone formation observed histologically. Statistical significance was determined using one-way ANOVA followed by Kruskal–Wallis multiple comparisons (***p < 0.001; ****p < 0.0001). Scale bars: 700 µm and 200 µm.

The pre-vascularized templates supported the development of significantly greater numbers of microvessels *in vivo*, as evidenced by Goldner trichrome staining (Fig. 4C) and confirmed by quantitative analysis (p < 0.0001; Fig. 4F). Despite increased vascularization, histological analysis revealed that the vascular channel templates supported significantly lower bone levels compared to the empty channel templates (p < 0.01; Fig. 4E). Overall, these histological analyses were consistent with the microCT findings, which demonstrated higher bone levels following implantation of the empty channel templates (Fig. 3C).

## 4. DISCUSSION

Bone tissue engineering aims to overcome the limitations of current treatments for large bone defect regeneration by developing grafts capable of supporting functional tissue regeneration. In this study, vascular microtissues fabricated using a co-culture of MSCs and HUVECs demonstrated the capacity to self-organize and form vascular networks when encapsulated in a fibrin hydrogel. A scaled-up graft was generated by integrating hypertrophic cartilage and vascular microtissues within a 3D-printed osteoconductive PCL framework coated with nano-hydroxyapatite, which supported cohesive fusion of the cartilage microtissues and the deposition of an extracellular matrix rich in sGAG and collagen. Bone formation was observed in both hypertrophic cartilage groups *in vivo*, however significantly greater levels of bone were observed in the empty channel group. In contrast, increased vascularization of the graft was observed following implantation of the pre-vascularized constructs. Collectively, these findings highlight, as proof of concept, the potential to combine microtissue-based strategies with advanced 3D biofabrication approaches to generate a hypertrophic, prevascularised template for bone repair.

Previous work has established microtissue and pellet-based strategies as promising approaches for directing endochondral bone formation, including hypertrophic cartilage microtissues that support vascularization and mineralization after implantation [ 31–35]. More recently, biofabrication approaches have used 3D-printed polymer frameworks to spatially organize cartilage microtissues and build larger, anatomically relevant tissues [36–42]. Building on these concepts, our study advances the bone tissue engineering field by integrating hypertrophic cartilage microtissues within an osteoconductive (nanoHA-coated) 3D printed PCL framework, enabling microtissue fusion within a millimetre-scale graft suitable for implantation. In terms of tissue quality, the extracellular matrix formed within the integrated constructs was consistent with reports showing that microtissue fusion and maturation can generate tissues rich in sGAG and collagen [43], and that combining cellular modules with supportive biomaterials, such as 3D printed polymers or macroporous hydrogels, can enhance scalable tissue formation [44–47]. To this end we also employed dynamic culture to mature these constructs, leveraging improved mass transport and mechanical cues [48–51] that have been widely associated with enhanced extracellular matrix deposition and more homogeneous tissue development in 3D scaffold-based systems.

Vascularization remains a central consideration for endochondral bone tissue engineering, because successful (re)modelling of a hypertrophic cartilage template depends on timely vascular invasion and nutrient delivery, particularly in larger constructs where diffusion limits rapidly emerge [52–53]. This has motivated prior work to prevascularize endochondral grafts or otherwise accelerate vessel ingrowth, although many approaches still rely primarily on simple seeding or encapsulation of endothelial (and support) cell suspensions within hydrogels or porous scaffolds [54–60]. In contrast, here we demonstrate a modular strategy in which vascular microtissues encapsulated in fibrin robustly sprout and self-organize microvascular structures *in vitro*, behavior in agreement with established endothelial spheroid sprouting paradigms in fibrin gels [61–63] and with bottom-up prevascularised spheroid systems for engineering other tissues [64–69]. Importantly, we further show that these vascular microtissues can be integrated alongside a hypertrophic cartilage template within a nanoHA-coated PCL framework, without impairing overall levels of extracellular matrix production. This modular “tissue-by-tissue” integration complements existing modular approaches [70–71], while offering a distinct advantage over strategies employing simple cell suspensions, where scalability and control of spatial organization can be more difficult to achieve.

The subcutaneous implantation in nude mouse model used here is an informative first-line assay to probe whether an engineered hypertrophic template can mineralize and initiate endochondral-like remodeling *in vivo* [72], largely independent of the confounding biomechanical and inflammatory cues present in orthotopic defects. In this setting, we demonstrated robust bone formation in both experimental groups by microCT and histology, yet significantly greater bone volume was observed in the empty-channel configuration, while the vascular-channel configuration supported significantly greater vascularization of the graft *in vivo*. These outcomes can be linked to the staged nature of endochondral ossification, where vascular invasion is a prerequisite that can potentially precede the mineralization of the cartilage template [73–77]. It should be noted that other ectopic studies report that pre-vascularization or scaffold architectural strategies can enhance bone formation in vivo [78–82], supporting the concept that vascular support can be beneficial, suggesting that outcomes observed in this model *in vivo* system likely depend on construct design and the timepoint of analysis. Endochondral bone formation in ectopic environments is a highly dynamic and temporally regulated process, characterized by an initial phase of cartilage formation and mineralization, followed by progressive remodeling in which cartilage is resorbed and replaced with bone, and where overall bone levels may stabilize or even decrease over time *in vivo* [83–85]. Contrary to our original hypothesis, pre - vascularization did not increase mineralized tissue formation at the 8-week timepoint, despite significantly enhancing vascular invasion. Instead, pre-vascularization is associated with increased vascular invasion and reduced residual cartilage, suggesting altered remodeling kinetics. The current study cannot determine whether the reduced bone volume reflects accelerated progression through the endochondral pathway or impaired bone formation. Additional *in vivo* timepoints would be required to address this question.

The reduced presence of cartilage observed within the pre-vascularized constructs *in vivo* is consistent with accelerated progression along the endochondral pathway and associated tissue (re)modelling and maturation. Channel-based architecture has previously been explored to accelerate vascularization and bone formation *in vivo* [31; 86–89], and it is plausible that the enhanced vascular infiltration observed here may have accelerated the transition from cartilage to bone and subsequent graft (re)modelling. Importantly, when endochondral approaches are evaluated in orthotopic critical-size defects, higher and more functional bone regeneration is often observed than in ectopic sites [84; 90–93], highlighting the contribution of the injury niche to regulating (re)modelling dynamics. Taken together, these findings suggest that the vascular-channel group may not be limited in osteogenic capacity but may instead reflect a shift in the temporal progression of endochondral (re)modelling, reinforcing the importance of selecting appropriate implantation models and timepoints when evaluating pre-vascularization strategies. An alternative explanation is that pre-vascularization in some way suppresses chondrogenesis, which in turn impacts endochondral bone formation *in vivo*, however no evidence of this was observed *in vitro*.

A key limitation of this study is that additional *in vitro* characterization could have further strengthened the mechanistic interpretation of the findings. In particular, mechanical testing of the scaled-up constructs (e.g., compressive and tensile properties) would have provided quantitative insight into functional maturation, while expanding molecular and immunohistochemistry to localize osteogenic, chondrogenic/hypertrophic and endothelial markers, could have clarified how microtissue interactions and channel design influenced the endochondral cascade. A second limitation is the use of an ectopic subcutaneous implantation model, which, while valuable for assessing vascularization and the bone forming potential of the graft in a controlled environment, does not recapitulate the inflammatory, mechanical, and biological cues of a healing fracture niche. Consequently, evaluation in a critical-size bone defect model would be an appropriate next step to directly assess the extent and functionality of bone regeneration and vascular integration at an orthotopic site.

In conclusion, this work demonstrates the successful biofabrication of vascular network–forming microtissues integrated within a 3D-printed, nanoHA-coated PCL framework for endochondral bone tissue engineering applications. The study successfully combined hypertrophic cartilage microtissues, vascular microtissues and osteoconductive 3D printed architectures, into a single integrated construct while maintaining tissue organization and biological function. Both the empty- and vascular-channel configurations supported bone formation following implantation, highlighting the capacity of these modular, self-organizing grafts to initiate endochondral bone tissue formation *in vivo*. Notably, the vascular-channel design promoted enhanced vascularization, supporting the promise of this vascular microtissue strategy for overcoming diffusion limitations and improving graft viability during early remodeling. Collectively, these findings position self-organized vascular microtissues combined with osteoinductive biofabrication platforms as a versatile route towards engineering scalable endochondral bone grafts, and future work will evaluate these designs in a rat critical-size bone defect model to assess regeneration within a more clinically relevant environment.

## REFERENCES

[1] Ho-Shui-Ling A, Bolander J, Rustom LE, Johnson AW, Luyten FP, Picart C. Bone regeneration strategies: Engineered scaffolds, bioactive molecules and stem cells current stage and future perspectives. Biomaterials. 2018 Oct; 180:143–162. doi: 10.1016/j.biomaterials.2018.07.017.

[2] Shi R, Huang Y, Ma C, Wu C, Tian W. Current advances for bone regeneration based on tissue engineering strategies. Front Med. 2019 Apr;13(2):160–188. doi: 10.1007/s11684-018-0629-9.

[3] Stanovici J, Le Nail LR, Brennan MA, Vidal L, Trichet V, Rosset P, Layrolle P. Bone regeneration strategies with bone marrow stromal cells in orthopaedic surgery. Curr Res Transl Med. 2016 Apr-Jun;64(2):83–90. doi: 10.1016/j.retram.2016.04.006.

[4] Majidinia M, Sadeghpour A, Yousefi B. The roles of signaling pathways in bone repair and regeneration. J Cell Physiol. 2018 Apr;233(4):2937–2948. doi: 10.1002/jcp.26042.

[5] Bharadwaz A, Jayasuriya AC. Recent trends in the application of widely used natural and synthetic polymer nanocomposites in bone tissue regeneration. Mater Sci Eng C Mater Biol Appl. 2020 May; 110:110698. doi: 10.1016/j.msec.2020.110698.

[6] Long F, Ornitz DM. Development of the endochondral skeleton. Cold Spring Harb Perspect Biol. 2013 Jan 1;5(1):a008334. doi: 10.1101/cshperspect.a008334.

[7] Krishnan L, Willett NJ, Guldberg RE. Vascularization strategies for bone regeneration. Ann Biomed Eng. 2014 Feb;42(2):432–44. doi: 10.1007/s10439-014-0969-9.

[8] Huang RL, Fu R, Yan Y, Liu C, Yang J, Xie Y, Li Q. Engineering hypertrophic cartilage grafts from lipoaspirate for critical-sized calvarial bone defect reconstruction: An adipose tissue-based developmental engineering approach. Bioeng Transl Med. 2022 Mar 24;7(3): e10312. doi: 10.1002/btm2.10312.

[9] Scotti C., Piccinini E., Takizawa H., Todorov A., Bourgine P., Papadimitropoulos A., et al. 2013. Engineering of a functional bone organ through endochondral ossification. Proc. Natl. Acad. Sci.U.S.A. doi: 110 3997–4002. 10.1073/pnas.1220108110.

[10] Matsiko A, Thompson EM, Lloyd-Griffith C, Cunniffe GM, Vinardell T, Gleeson JP, Kelly DJ, O’Brien FJ. An endochondral ossification approach to early stage bone repair: Use of tissue-engineered hypertrophic cartilage constructs as primordial templates for weight-bearing bone repair. J Tissue Eng Regen Med. 2018 Apr;12(4):e2147–e2150. doi: 10.1002/term.2638.

[11] Fröhlich, M., Grayson, W., Marolt, D., Gimble, J., Kregar-Velikonja, N., Vunjak-Novakovic, G. Bone grafts engineered from human adipose-derived stem cells in perfusion bioreactor culture. Tissue Eng Part A. 2009, 16;16(1):179–189. doi: 10.1089/ten.tea.2009.0164.

[12] de Silva LS, Kuijpers CJ, Van Cann EM, Rosenberg AJWP, van Es RJJ, Gawlitta D. The impact of vascular supply on endochondral bone regeneration in centimeter-sized porous chambers. Tissue Eng Part A. 2026 Jan;32(1-2):21–30. doi: 10.1089/ten.tea.2025.0045.

[13] Bai L, Zhou D, Li G, Liu J, Chen X, Su J. Engineering bone/cartilage organoids: strategy, progress, and application. Bone Res. 2024 Nov 20;12(1):66. doi: 10.1038/s41413-024-00376-y.

[14] Peng L, Decoene I, Svitina H, Papantoniou I. Synchronizing the Osteochondral Regeneration Process through Spatial Patterning of Stable and Hypertrophic Cartilage Organoids. Adv Mater. 2026 May 8: e16189. doi: 10.1002/adma.202516189.

[15] Albillos Sanchez A, Marks MP, Casademunt P, Seijas-Gamardo A, Papantoniou I, Moroni L, Mota C. Packed for Ossification: High-Density Bioprinting of hPDC Spheroids in HAMA Toward Endochondral Ossification. Adv Healthc Mater. 2026 Mar 15: e05855. doi: 10.1002/adhm.202505855.

[16] Achilli TM, Meyer J, Morgan JR. Advances in the formation, use and understanding of multi-cellular spheroids. Expert Opin Biol Ther. 2012 Oct;12(10):1347–60. doi: 10.1517/14712598.2012.707181.

[17] Burdis R, Kelly DJ. Biofabrication and bioprinting using cellular aggregates, microtissues and organoids for the engineering of musculoskeletal tissues. Acta Biomater. 2021 May; 126:1–14. doi: 10.1016/j.actbio.2021.03.016.

[18] F. C. Teixeira, V. Joris, M. van Griensven, L. Moroni, and C. Mota. Pre-Vascularized hMSC and hPDC Spheroids as Building Block Units for Bone Tissue Engineering. Advanced Materials Interfaces13, no. 2 (2026): e00804. doi: 10.1002/admi.202500804.

[19] Wrublewsky S, Schultz J, Ammo T, Bickelmann C, Metzger W, Später T, Pohlemann T, Menger MD, Laschke MW. Biofabrication of prevascularized spheroids for bone tissue engineering by fusion of microvascular fragments with osteoblasts. Front Bioeng Biotechnol. 2024 Sep 10; 12:1436519. doi: 10.3389/fbioe.2024.1436519.

[20] Tiruvannamalai Annamalai R, Rioja AY, Putnam AJ, Stegemann JP. Vascular Network Formation by Human Microvascular Endothelial Cells in Modular Fibrin Microtissues. ACS Biomater Sci Eng. 2016 Nov 14;2(11):1914–1925. doi: 10.1021/acsbiomaterials.6b00274.

[21] Ohori-Morita Y, Niibe K, Limraksasin P, Nattasit P, Miao X, Yamada M, Mabuchi Y, Matsuzaki Y, Egusa H. Novel Mesenchymal Stem Cell Spheroids with Enhanced Stem Cell Characteristics and Bone Regeneration Ability. Stem Cells Transl Med. 2022.

[22] Lee J, Seok JM, Huh SJ, Byun H, Lee S, Park SA, Shin H. 3D printed micro-chambers carrying stem cell spheroids and pro-proliferative growth factors for bone tissue regeneration. Biofabrication. 2020 Dec 17;13(1). doi: 10.1088/1758-5090/abc39c.

[23] Shanbhag S, Suliman S, Mohamed-Ahmed S, Kampleitner C, Hassan MN, Heimel P, Dobsak T, Tangl S, Bolstad AI, Mustafa K. Bone regeneration in rat calvarial defects using dissociated or spheroid mesenchymal stromal cells in scaffold-hydrogel constructs. Stem Cell Res Ther. 2021 Nov 14;12(1):575. doi: 10.1186/s13287-021-02642-w.

[24] Yamada Y, Okano T, Orita K, Makino T, Shima F, Nakamura H. 3D-cultured small size adipose-derived stem cell spheroids promote bone regeneration in the critical-sized bone defect rat model. Biochem Biophys Res Commun. 2022 May 7; 603:57–62. doi: 10.1016/j.bbrc.2022.03.027.

[25] Prabhakaran V, Melchels FPW, Murray LM, Paxton JZ. Engineering three-dimensional bone macro-tissues by guided fusion of cell spheroids. Front Endocrinol (Lausanne). 2023 Dec 19; 14:1308604. doi: 10.3389/fendo.2023.1308604.

[26] Chae Y, Jang K, Lee S, Kim YH, Jin S, Shim KM, Kang SS, Kim SE. Bone regeneration using adipose derived stem cell spheroids within 3D printed scaffolds in a rabbit radial defect model. Sci Rep. 2025 Dec 16;15(1):43954. doi: 10.1038/s41598-025-25581-5.

[27] Fang Y, Ji M, Wu B, Xu X, Wang G, Zhang Y, Xia Y, Li Z, Zhang T, Sun W, Xiong Z. Engineering Highly Vascularized Bone Tissues by 3D Bioprinting of Granular Prevascularized Spheroids. ACS Appl Mater Interfaces. 2023 Sep 20;15(37):43492–43502. doi: 10.1021/acsami.3c08550.

[28] Kim W, Jang CH, Kim G. Bone tissue engineering supported by bioprinted cell constructs with endothelial cell spheroids. Theranostics. 2022 Jul 11;12(12):5404–5417. doi: 10.7150/thno.74852.

[29] Nulty J, Burdis R, Kelly DJ. Biofabrication of Prevascularised Hypertrophic Cartilage Microtissues for Bone Tissue Engineering. Front Bioeng Biotechnol. 2021 Jun 7; 9:661989. doi: 10.3389/fbioe.2021.661989.

[30] Eichholz KF, Von Euw S, Burdis R, Kelly DJ, Hoey DA. Development of a New Bone-Mimetic Surface Treatment Platform: Nanoneedle Hydroxyapatite (nnHA) Coating. Adv Healthc Mater. 2020 Dec;9(24): e2001102. doi: 10.1002/adhm.202001102.

[31] Sheehy EJ, Vinardell T, Toner ME, Buckley CT, Kelly DJ. Altering the architecture of tissue engineered hypertrophic cartilaginous grafts facilitates vascularisation and accelerates mineralisation. PLoS One. 2014 Mar 4;9(3): e90716. doi: 10.1371/journal.pone.0090716.

[32] Sheehy EJ, Kelly DJ, O’Brien FJ. Biomaterial-based endochondral bone regeneration: a shift from traditional tissue engineering paradigms to developmentally inspired strategies. Mater Today Bio. 2019 May 31; 3:100009. doi: 10.1016/j.mtbio.2019.100009.

[33] Bernhard, Jonathan C., Darja Marolt Presen, Ming Li, Xavier Monforte, James Ferguson, Gabriele Leinfellner, Patrick Heimel, Susanna L. Betti, Sharon Shu, Andreas H. Teuschl-Woller, and, et al. 2022. Effects of Endochondral and Intramembranous Ossification Pathways on Bone Tissue Formation and Vascularization in Human Tissue-Engineered Grafts. Cells 11, no. 19: 3070. 10.3390/cells11193070.

[34] Huang RL, Fu R, Yan Y, Liu C, Yang J, Xie Y, Li Q. Engineering hypertrophic cartilage grafts from lipoaspirate for critical-sized calvarial bone defect reconstruction: An adipose tissue-based developmental engineering approach. Bioeng Transl Med. 2022 Mar 24;7(3): e10312. doi: 10.1002/btm2.10312.

[35] Schott NG, Kaur G, Coleman RM, Stegemann JP. Modular, Vascularized Hypertrophic Cartilage Constructs for Bone Tissue Engineering Applications. Tissue Eng Part A. 2025 Dec;31(23-24):1297–1308. doi: 10.1089/ten.tea.2024.0367.

[36] Daly AC, Kelly DJ. Biofabrication of spatially organised tissues by directing the growth of cellular spheroids within 3D printed polymeric microchambers. Biomaterials. 2019 Mar; 197:194–206. doi: 10.1016/j.biomaterials.2018.12.028.

[37] Lee J, Seok JM, Huh SJ, Byun H, Lee S, Park SA, Shin H. 3D printed micro-chambers carrying stem cell spheroids and pro-proliferative growth factors for bone tissue regeneration. Biofabrication. 2020 Dec 17;13(1). doi: 10.1088/1758-5090/abc39c.

[38] Ahmad T, Byun H, Lee J, Madhurakat Perikamana SK, Shin YM, Kim EM, Shin H. Stem cell spheroids incorporating fibers coated with adenosine and polydopamine as a modular building block for bone tissue engineering. Biomaterials. 2020 Feb; 230:119652. doi: 10.1016/j.biomaterials.2019.119652.

[39] Yamada Y, Okano T, Orita K, Makino T, Shima F, Nakamura H. 3D-cultured small size adipose-derived stem cell spheroids promote bone regeneration in the critical-sized bone defect rat model. Biochem Biophys Res Commun. 2022 May 7; 603:57–62. doi: 10.1016/j.bbrc.2022.03.027.

[40] Liu X, Li L, Gaihre B, Park S, Li Y, Terzic A, Elder BD, Lu L. Scaffold-Free Spheroids with Two-Dimensional Heteronano-Layers (2DHNL) Enabling Stem Cell and Osteogenic Factor Codelivery for Bone Repair. ACS Nano. 2022 Feb 22;16(2):2741–2755. doi: 10.1021/acsnano.1c09688.

[41] Hall GN, Chandrakar A, Pastore A, Ioannidis K, Moisley K, Cirstea M, Geris L, Moroni L, Luyten FP, Wieringa P, Papantoniou I. Engineering bone-forming biohybrid sheets through the integration of melt electrowritten membranes and cartilaginous micro spheroids. Acta Biomater. 2023 Jul 15; 165:111–124. doi: 10.1016/j.actbio.2022.10.037.

[42] Chae Y, Jang K, Lee S, Kim YH, Jin S, Shim KM, Kang SS, Kim SE. Bone regeneration using adipose derived stem cell spheroids within 3D printed scaffolds in a rabbit radial defect model. Sci Rep. 2025 Dec 16;15(1):43954. doi: 10.1038/s41598-025-25581-5.

[43] Spagnuolo FD, Kronemberger GS, Storey KJ, Kelly DJ. The maturation state and density of human cartilage microtissues influence their fusion and development into scaled-up grafts. Acta Biomater. 2025 Mar 1; 194:109–121. doi: 10.1016/j.actbio.2025.01.024.

[44] Fan C, Wang DA. Macroporous Hydrogel Scaffolds for Three-Dimensional Cell Culture and Tissue Engineering. Tissue Eng Part B Rev. 2017 Oct;23(5):451–461. doi: 10.1089/ten.TEB.2016.0465.

[45] Li L, Wu W, Zhu Q, Peng M, Dong Y, Liu H, Zhang J, Gou M. Advanced cell-adaptable hydrogels for bioprinting. Bioact Mater. 2025 Aug 6; 53:831–854. doi: 10.1016/j.bioactmat.2025.07.044.

[46] Khan, A., Gholap, A., Grewal, N., Jun, Z., Khalid, M., Zhang, H-J. Advances in smart hybrid scaffolds: A strategic approach for regenerative clinical applications, Engineered Regeneration, Volume 6, 2025, Pages 85-110, doi: 10.1016/j.engreg.2025.02.002.

[47] Puiggalí-Jou A, Hui IB, Fernandez-Rico C, Zenobi-Wong M. The Space Within: How Architected Voids Promote Tissue Formation. Adv Mater. 2026 Feb;38(7): e07385. doi: 10.1002/adma.202507385. Epub 2025 Nov 28. PMID: 41312612.

[48] Daly AC, Sathy BN, Kelly DJ. Engineering large cartilage tissues using dynamic bioreactor culture at defined oxygen conditions. J Tissue Eng. 2018 Jan 24; 9:2041731417753718. doi: 10.1177/2041731417753718.

[49] Kronstadt SM, Patel DB, Born LJ, Levy D, Lerman MJ, Mahadik B, McLoughlin ST, Fasuyi A, Fowlkes L, Van Heyningen LH, Aranda A, Abadchi SN, Chang KH, Hsu ATW, Bengali S, Harmon JW, Fisher JP, Jay SM. Mesenchymal Stem Cell Culture within Perfusion Bioreactors Incorporating 3D-Printed Scaffolds Enables Improved Extracellular Vesicle Yield with Preserved Bioactivity. Adv Healthc Mater. 2023 Aug;12(20): e2300584. doi: 10.1002/adhm.202300584.

[50] Wu G, Chen S, Li Q, Zhang M, Cao F, Wei J, Guo L, Li P, Wei X, Zhang Q. Mechanical cues enhance chondrocyte function: Insights from mechanoreception, regulation, and biological responses. Mechanobiol Med. 2025 Nov 19;3(4):100164. doi: 10.1016/j.mbm.2025.100164.

[51] Chariyev-Prinz F, Burdis R, Kelly DJ. Chondrogenic Maturation Governs hMSC Mechanoresponsiveness to Dynamic Compression. Bioengineering (Basel). 2025 Oct 3;12(10):1075. doi: 10.3390/bioengineering12101075.

[52] Lopes D, Martins-Cruz C, Oliveira MB, Mano JF. Bone physiology as inspiration for tissue regenerative therapies. Biomaterials. 2018 Dec; 185:240–275. doi: 10.1016/j.biomaterials.2018.09.028.

[53] Schott NG, Kaur G, Coleman RM, Stegemann JP. Modular, Vascularized Hypertrophic Cartilage Constructs for Bone Tissue Engineering Applications. Tissue Eng Part A. 2025 Dec;31(23-24):1297–1308. doi: 10.1089/ten.tea.2024.0367.

[54] Farrell E, Both SK, Odörfer KI, Koevoet W, Kops N, O’Brien FJ, Baatenburg de Jong RJ, Verhaar JA, Cuijpers V, Jansen J, Erben RG, van Osch GJ. In-vivo generation of bone via endochondral ossification by in-vitro chondrogenic priming of adult human and rat mesenchymal stem cells. BMC Musculoskelet Disord. 2011 Jan 31; 12:31. doi: 10.1186/1471-2474-12-31.

[55] Yang W, Both SK, van Osch GJ, Wang Y, Jansen JA, Yang F. Performance of different three-dimensional scaffolds for in vivo endochondral bone generation. Eur Cell Mater. 2014 Jun 10; 27:350–64. doi: 10.22203/ecm. v027a25.

[56] Sheehy EJ, Vinardell T, Toner ME, Buckley CT, Kelly DJ. Altering the architecture of tissue engineered hypertrophic cartilaginous grafts facilitates vascularisation and accelerates mineralisation. PLoS One. 2014 Mar 4;9(3): e90716. doi: 10.1371/journal.pone.0090716.

[57] Yan Y, Chen H, Zhang H, Guo C, Yang K, Chen K, Cheng R, Qian N, Sandler N, Zhang YS, Shen H, Qi J, Cui W, Deng L. Vascularized 3D printed scaffolds for promoting bone regeneration. Biomaterials. 2019 Jan;190–191:97-110. doi: 10.1016/j.biomaterials.2018.10.033.

[58] Chiesa I, De Maria C, Lapomarda A, Fortunato GM, Montemurro F, Di Gesù R, Tuan RS, Vozzi G, Gottardi R. Endothelial cells support osteogenesis in an in vitro vascularized bone model developed by 3D bioprinting. Biofabrication. 2020 Feb 19;12(2):025013. doi: 10.1088/1758-5090/ab6a1d.

[59] Mikael PE, Golebiowska AA, Xin X, Rowe DW, Nukavarapu SP. Evaluation of an Engineered Hybrid Matrix for Bone Regeneration via Endochondral Ossification. Ann Biomed Eng. 2020 Mar;48(3):992–1005. doi: 10.1007/s10439-019-02279-0.

[60] Grottkau BE, Hui Z, Ran C, Pang Y. Fabricating vascularized, anatomically accurate bone grafts using 3D bioprinted sectional bone modules, in-situ angiogenesis, BMP-2 controlled release, and bioassembly. Biofabrication. 2024 Jul 16;16(4). doi: 10.1088/1758-5090/ad5f56.

[61] Murphy KC, Fang SY, Leach JK. Human mesenchymal stem cell spheroids in fibrin hydrogels exhibit improved cell survival and potential for bone healing. Cell Tissue Res. 2014 Jul;357(1):91–9. doi: 10.1007/s00441-014-1830-z.

[62] Tiruvannamalai Annamalai R, Rioja AY, Putnam AJ, Stegemann JP. Vascular Network Formation by Human Microvascular Endothelial Cells in Modular Fibrin Microtissues. ACS Biomater Sci Eng. 2016 Nov 14;2(11):1914–1925. doi: 10.1021/acsbiomaterials.6b00274.

[63] Heo DN, Hospodiuk M, Ozbolat IT. Synergistic interplay between human MSCs and HUVECs in 3D spheroids laden in collagen/fibrin hydrogels for bone tissue engineering. Acta Biomater. 2019 Sep 1; 95:348–356. doi: 10.1016/j.actbio.2019.02.046.

[64] Kanczler JM, Wells JA, Oreffo ROC. Endothelial Cells: Co-culture Spheroids. Methods Mol Biol. 2021; 2206:47–56. doi: 10.1007/978-1-0716-0916-3_5.

[65] Rahimnejad M, Nasrollahi Boroujeni N, Jahangiri S, Rabiee N, Rabiee M, Makvandi P, Akhavan O, Varma RS. Prevascularized Micro-/Nano-Sized Spheroid/Bead Aggregates for Vascular Tissue Engineering. Nanomicro Lett. 2021 Aug 18;13(1):182. doi: 10.1007/s40820-021-00697-1.

[66] Kook MG, Lee SE, Shin N, Kong D, Kim DH, Kim MS, Kang HK, Choi SW, Kang KS. Generation of Cortical Brain Organoid with Vascularization by Assembling with Vascular Spheroid. Int J Stem Cells. 2022 Feb 28;15(1):85–94. doi: 10.15283/ijsc21157.

[67] Philippon EML, van Rooijen LJE, Khodadust F, van Hamburg JP, van der Laken CJ, Tas SW. A novel 3D spheroid model of rheumatoid arthritis synovial tissue incorporating fibroblasts, endothelial cells, and macrophages. Front Immunol. 2023 Jul 20; 14:1188835. doi: 10.3389/fimmu.2023.1188835.

[68] Escudero M, Vaysse L, Eke G, Peyrou M, Villarroya F, Bonnel S, Jeanson Y, Boyer L, Vieu C, Chaput B, Yao X, Deschaseaux F, Parny M, Raymond-Letron I, Dani C, Carrière A, Malaquin L, Casteilla L. Scalable Generation of Pre-Vascularized and Functional Human Beige Adipose Organoids. Adv Sci (Weinh). 2023 Nov;10(31): e2301499. doi: 10.1002/advs.202301499.

[69] Miao Y, Pek NM, Tan C, Jiang C, Yu Z, Iwasawa K, Shi M, Kechele DO, Sundaram N, Pastrana-Gomez V, Sinner DI, Liu X, Lin KC, Na CL, Kishimoto K, Yang MC, Maharjan S, Tchieu J, Whitsett JA, Zhang YS, McCracken KW, Rottier RJ, Kotton DN, Helmrath MA, Wells JM, Takebe T, Zorn AM, Chen YW, Guo M, Gu M. Co-development of mesoderm and endoderm enables organotypic vascularization in lung and gut organoids. Cell. 2025 Aug 7;188(16):4295–4313.e27. doi: 10.1016/j.cell.2025.05.041.

[70] Nulty J, Burdis R, Kelly DJ. Biofabrication of Prevascularised Hypertrophic Cartilage Microtissues for Bone Tissue Engineering. Front Bioeng Biotechnol. 2021 Jun 7; 9:661989. doi: 10.3389/fbioe.2021.661989.

[71] Schott NG, Kaur G, Coleman RM, Stegemann JP. Modular, Vascularized Hypertrophic Cartilage Constructs for Bone Tissue Engineering Applications. Tissue Eng Part A. 2025 Dec;31(23-24):1297–1308. doi: 10.1089/ten.tea.2024.0367.

[72] Ng J, Wei Y, Zhou B, Bhumiratana S, Burapachaisri A, Guo E, Vunjak-Novakovic G. Ectopic implantation of juvenile osteochondral tissues recapitulates endochondral ossification. J Tissue Eng Regen Med. 2018 Feb;12(2):468–478. doi: 10.1002/term.2500.

[73] van der Pluijm G, Deckers M, Sijmons B, de Groot H, Bird J, Wills R, Papapoulos S, Baxter A, Löwik C. In vitro and in vivo endochondral bone formation models allow identification of anti-angiogenic compounds. Am J Pathol. 2003 Jul;163(1):157–63. doi: 10.1016/S0002-9440(10)63639-5.

[74] Mark H, Penington A, Nannmark U, Morrison W, Messina A. Microvascular invasion during endochondral ossification in experimental fractures in rats. Bone. 2004 Aug;35(2):535–42. doi: 10.1016/j.bone.2004.04.010.

[75] Grosso A, Burger MG, Lunger A, Schaefer DJ, Banfi A, Di Maggio N. It Takes Two to Tango: Coupling of Angiogenesis and Osteogenesis for Bone Regeneration. Front Bioeng Biotechnol. 2017 Nov 3; 5:68. doi: 10.3389/fbioe.2017.00068.

[76] Bernhard JC, Marolt Presen D, Li M, Monforte X, Ferguson J, Leinfellner G, Heimel P, Betti SL, Shu S, Teuschl-Woller AH, Tangl S, Redl H, Vunjak-Novakovic G. Effects of Endochondral and Intramembranous Ossification Pathways on Bone Tissue Formation and Vascularization in Human Tissue-Engineered Grafts. Cells. 2022 Sep 29;11(19):3070. doi: 10.3390/cells11193070.

[77] Zhang X, Jiang W, Wu X, Xie C, Zhang Y, Li L, Gu Y, Hu Z, Zhai X, Liang R, Zhang T, Sun W, Ye J, Wei W, Wang X, Hong Y, Zhang S, Cai Y, Zou X, Hu Y, Ouyang H. Divide-and-conquer strategy with engineered ossification center organoids for rapid bone healing through developmental cell recruitment. Nat Commun. 2025 Jul 4;16(1):6200. doi: 10.1038/s41467-025-61619-y.

[78] Sheehy EJ, Vinardell T, Toner ME, Buckley CT, Kelly DJ. Altering the architecture of tissue engineered hypertrophic cartilaginous grafts facilitates vascularisation and accelerates mineralisation. PLoS One. 2014 Mar 4;9(3): e90716. doi: 10.1371/journal.pone.0090716.

[79] García, J.R., García, A.J. Biomaterial-mediated strategies targeting vascularization for bone repair. Drug Deliv. and Transl. Res. 6, 77–95 (2016). 10.1007/s13346-015-0236-0.

[80] Nulty J, Freeman FE, Browe DC, Burdis R, Ahern DP, Pitacco P, Lee YB, Alsberg E, Kelly DJ. 3D bioprinting of prevascularised implants for the repair of critically-sized bone defects. Acta Biomater. 2021 May; 126:154–169. doi: 10.1016/j.actbio.2021.03.003.

[81] Bhatia S, Hipwood L, Claxton B, Bessot A, Weekes A, Sokolowski K, Mashimo T, Bock N, McGovern J. Divergent effects of premineralization and prevascularization on osteogenesis and vascular integration in humanized tissue engineered bone constructs. Acta Biomater. 2025 Jul 1; 201:665–683. doi: 10.1016/j.actbio.2025.06.019.

[82] Dazzi C, Eichholz KF, Freeman FE, Kelly DJ, Checa S. An in silico study reveals how architectural and mechanical cues jointly regulate angiogenesis and bone regeneration in 3D printed scaffolds. Comput Biol Med. 2025 Sep; 195:110574. doi: 10.1016/j.compbiomed.2025.110574.

[83] Thompson EM, Matsiko A, Farrell E, Kelly DJ, O’Brien FJ. Recapitulating endochondral ossification: a promising route to in vivo bone regeneration. J Tissue Eng Regen Med. 2015 Aug;9(8):889–902. doi: 10.1002/term.1918.

[84] Freeman FE, Brennan MÁ, Browe DC, Renaud A, De Lima J, Kelly DJ, McNamara LM, Layrolle P. A. Developmental Engineering-Based Approach to Bone Repair: Endochondral Priming Enhances Vascularization and New Bone Formation in a Critical Size Defect. Front Bioeng Biotechnol. 2020 Mar 31; 8:230. doi: 10.3389/fbioe.2020.00230.

[85] Acevedo Rua L, Mumme M, Manferdini C, Darwiche S, Khalil A, Hilpert M, Buchner DA, Lisignoli G, Occhetta P, von Rechenberg B, Haug M, Schaefer DJ, Jakob M, Caplan A, Martin I, Barbero A, Pelttari K. Engineered nasal cartilage for the repair of osteoarthritic knee cartilage defects. Sci Transl Med. 2021 Sep;13(609): eaaz4499. doi: 10.1126/scitranslmed.aaz4499.

[86] Byambaa B, Annabi N, Yue K, Trujillo-de Santiago G, Alvarez MM, Jia W, Kazemzadeh-Narbat M, Shin SR, Tamayol A, Khademhosseini A. Bioprinted Osteogenic and Vasculogenic Patterns for Engineering 3D Bone Tissue. Adv Healthc Mater. 2017 Aug;6(16):10.1002/adhm.201700015. doi: 10.1002/adhm.201700015.

[87] Klotz BJ, Lim KS, Chang YX, Soliman BG, Pennings I, Melchels FPW, Woodfield TBF, Rosenberg AJ, Malda J, Gawlitta D. Engineering of a complex bone tissue model with Endothelialised channels and capillary-like networks. Eur Cell Mater. 2018 May 30; 35:335–348. doi: 10.22203/eCM.v035a23.

[88] Shen M, Wang L, Gao Y, Feng L, Xu C, Li S, Wang X, Wu Y, Guo Y, Pei G. 3D bioprinting of in situ vascularized tissue engineered bone for repairing large segmental bone defects. Mater Today Bio. 2022 Aug 8; 16:100382. doi: 10.1016/j.mtbio.2022.100382.

[89] Ghobadi E, Yahay Z, Nouri N, Karamali F, Masaeli E. 3D printing of an anatomically shaped bone model inspired by vascularized tubular bone structure. Biomater Adv. 2025 Nov; 176:214348. doi: 10.1016/j.bioadv.2025.214348.

[90] Dang PN, Herberg S, Varghai D, Riazi H, Varghai D, McMillan A, Awadallah A, Phillips LM, Jeon O, Nguyen MK, Dwivedi N, Yu X, Murphy WL, Alsberg E. Endochondral Ossification in Critical-Sized Bone Defects via Readily Implantable Scaffold-Free Stem Cell Constructs. Stem Cells Transl Med. 2017 Jul;6(7):1644–1659. doi: 10.1002/sctm.16-0222.

[91] Tam WL, Freitas Mendes L, Chen X, Lesage R, Van Hoven I, Leysen E, Kerckhofs G, Bosmans K, Chai YC, Yamashita A, Tsumaki N, Geris L, Roberts SJ, Luyten FP. Human pluripotent stem cell-derived cartilaginous organoids promote scaffold-free healing of critical size long bone defects. Stem Cell Res Ther. 2021 Sep 25;12(1):513. doi: 10.1186/s13287-021-02580-7.

[92] Longoni A, Utomo L, Robinson A, Levato R, Rosenberg AJWP, Gawlitta D. Acceleration of Bone Regeneration Induced by a Soft-Callus Mimetic Material. Adv Sci (Weinh). 2022 Feb;9(6): e2103284. doi: 10.1002/advs.202103284.

[93] Garcia Garcia A, Prithiviraj S, Raina DB, Schmidt T, Gonzalez Anton S, Rabanal Cajal L, Hidalgo Gil D, Tägil M, Hyrenius-Wittsten A, Dahlgren MW, Kahn R, Bourgine PE. Engineered and decellularized human cartilage graft exhibits intrinsic immunosuppressive properties and full skeletal repair capacity. Proc Natl Acad Sci U S A. 2026 Jan 13;123(2): e2507185123. doi: 10.1073/pnas.2507185123.

