## Supplementary figures for "Integrating vascular and hypertrophic cartilage microtissues to fabricate scaled-up grafts for endochondral bone tissue engineering"

**Supplementary material**

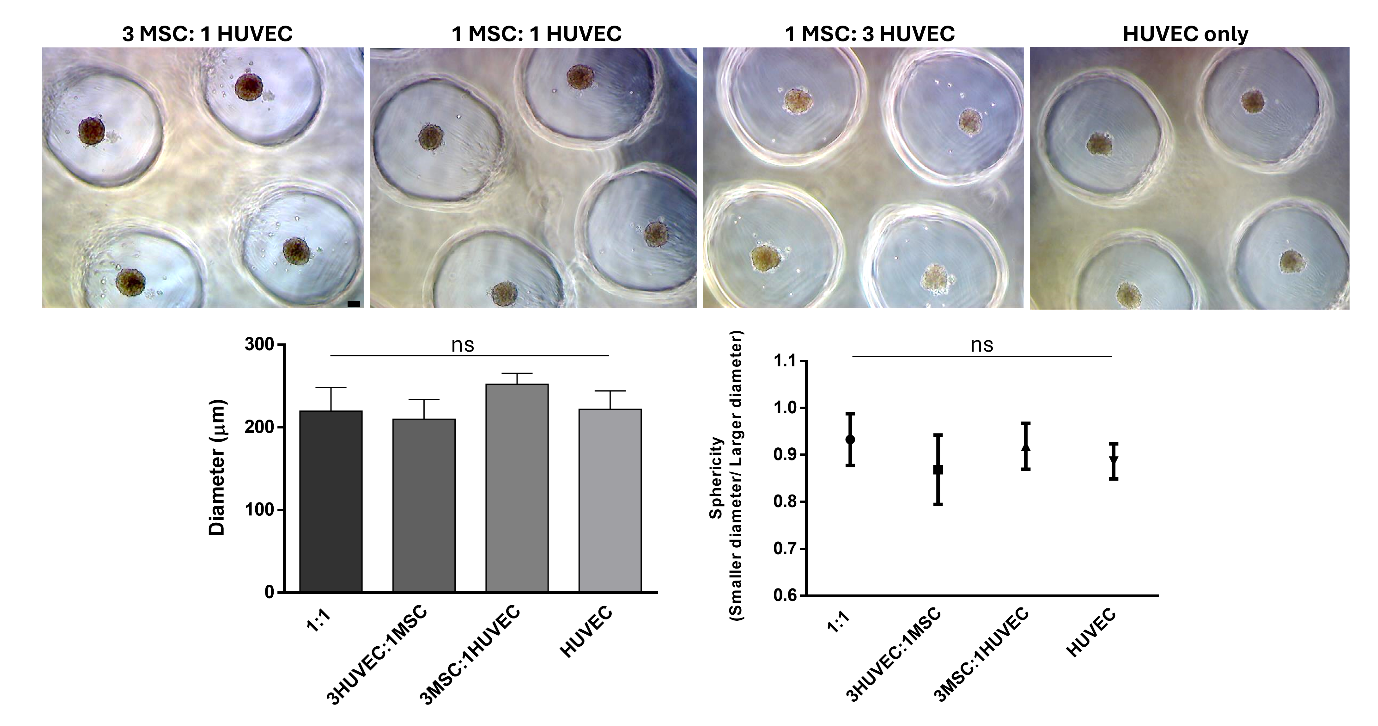

**Figure 1: Mean values of the larger and smaller diameters, as well as sphericity values, of co-culture spheroids composed of human BM-MSCs and HUVECs at different cell ratios.** Representative phase-contrast micrographs of co-culture spheroids after 24 h of culture are shown for the following cell ratios: 3 MSC:1 HUVEC (A), 1 MSC:1 HUVEC (B), 1 MSC:3 HUVEC (C), and HUVEC only (D). (E) Mean values of the major and minor diameters of co-culture spheroids according to cell ratios after 24 h of culture. (F) Sphericity values based on spheroid diameters according to cell ratios after 24 h of culture. Abbreviation: ns – not statistically significant. Scale bar: 100 μm.

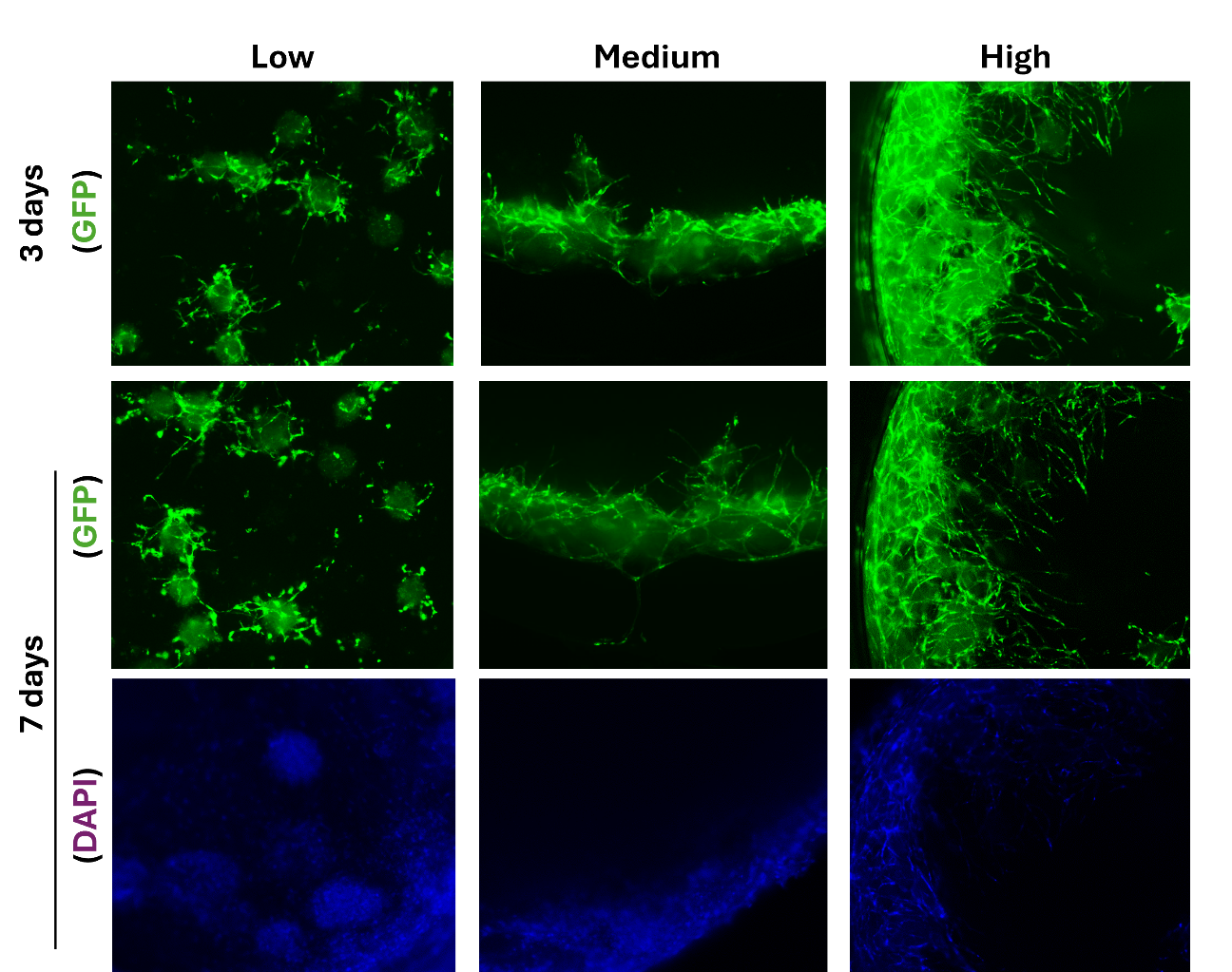

**Figure 2: Fluorescence images of vascular spheroids at a 3 MSC:1 HUVEC ratio seeded at different densities in fibrin-based bioink.** Representative images after 3 days of culture show spheroids seeded at low (A), medium (B), and high (C) densities. Note the presence of branching and the onset of sprouting in spheroids seeded at medium density (B). Increased branching and elongation are observed at high seeding density (C, arrow). Fluorescence micrographs after 7 days of culture show spheroids seeded at low (D), medium (E), and high (F) densities. Note the elongation of branches at medium density (E) and more pronounced elongation at high density (F). Fluorescence micrographs with DAPI staining of vascular spheroids seeded at low (G), medium (H), and high (I) densities. Abbreviation: GFP – green fluorescent protein.

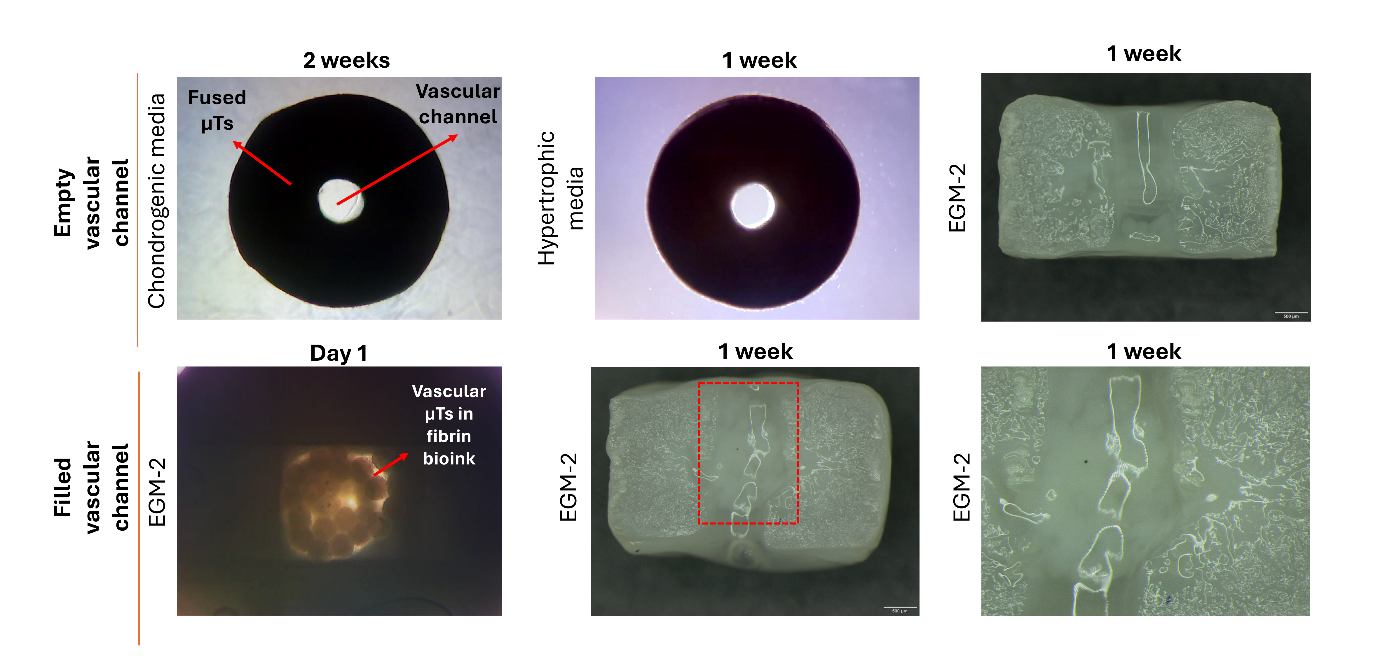

**Figure 3: Seeding of hypertrophic and vascular spheroids into a 3D-printed PCL scaffold.** Phase-contrast micrograph of fully fused spheroids within the 3D-printed PCL scaffold after 2 weeks of chondrogenic induction. Note the empty vascular channel. Phase-contrast micrograph of fully fused spheroids within the 3D-printed PCL scaffold after 1 week in hypertrophic induction medium. Note that the spheroids are adhered to the scaffold. Macroscopic image of the construct after 1 week in EGM-2 medium. Vascular spheroids seeded in fibrin bioink within the central channel of the scaffold after 1 day of culture. Macroscopic image of vascular spheroids after 1 week of culture in EGM-2 medium. The red rectangle indicates the central channel containing the seeded spheroids. Magnified view of vascular spheroids within the central channel of the PCL scaffold after 1 week in EGM-2 medium. Abbreviation: EGM-2 – endothelial cell growth medium-2. Scale bar: 500 μm.

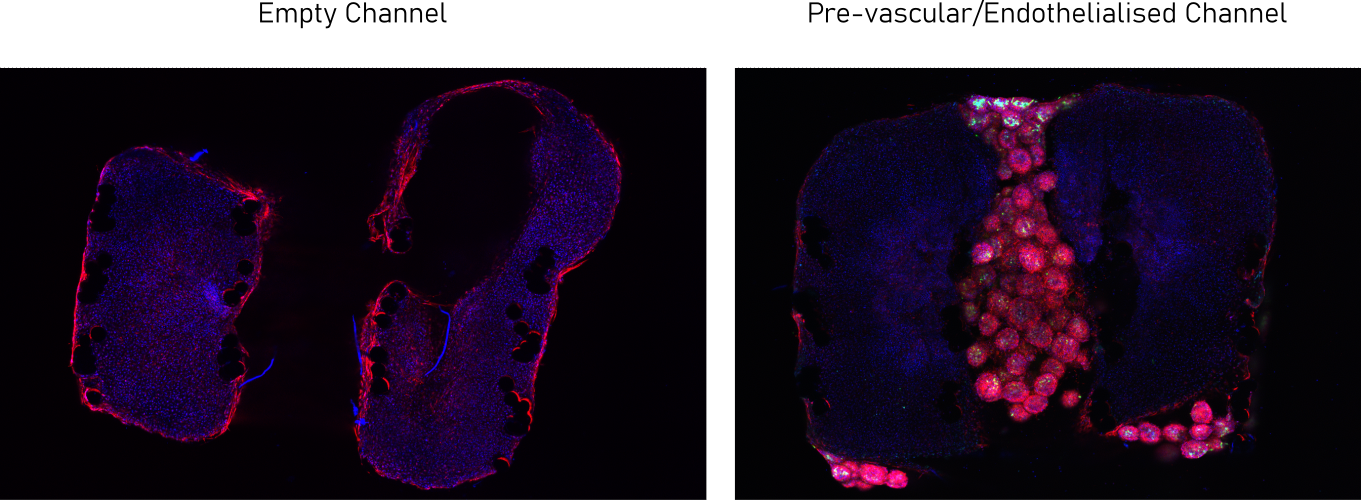

**Figure 4: Fluorescence micrographs of constructs with an empty and pre-vascular channel, maintained in chondrogenic induction medium for 2 weeks followed by hypertrophic induction medium for 1 week.**
